# Neural Correlates of Listening States, Cognitive Load, and Selective Attention in an Ecological Multi-Talker Scenario

**DOI:** 10.64898/2026.03.13.711289

**Authors:** Payam Shahsavari Baboukani, Rodrigo Ordoñez, Carina Gravesen, Jan Østergaard, Mike Lind Rank, Emina Aličković, Alvaro Fuentes Cabrera

## Abstract

This study assessed neural responses to continuous speech to classify listening state, cognitive load, and selective auditory attention in complex acoustic environments. EEG was recorded while participants listened to concurrent male and female talkers under two conditions: active listening, where attention was directed to one of two competing speakers (target vs. masker), or passive listening, where attention was diverted to a visual task. Cognitive load was varied by manipulating target-to-masker (TMR) ratio (TMR: +7 dB, −7 dB), with lower TMR representing more demanding listening conditions. Spectral EEG features across frequency bands were ranked with univariate statistics and used to classify listening state (active vs passive) and cognitive load (low vs. high TMR). Auditory attention decoding (AAD) was performed using linear stimulus reconstruction to identify the target talker during active listening. Classification of listening state achieved 90.3% accuracy, and AAD reached 84.4% accuracy, demonstrating robust tracking of attentional engagement. In contrast, classification of cognitive load was near chance, suggesting that more extreme acoustic manipulations may be required to elicit distinct neural signatures. Comparable performance using a reduced set of electrodes near the ear indicates the potential for integration with wearable hearing devices. Overall, these results demonstrate that EEG can distinguish attentional states and selectively track target speech in realistic auditory scenarios. The findings provide a foundation for future applications in monitoring listening behavior, supporting auditory processing, and improving brain-controlled hearing aids in complex acoustic environments.

**Highlights:**

- Listening state (active vs. passive) can be classified from EEG spectral features.
- Attended speech can be decoded by reconstructing speech envelopes from EEG.
- Comparable accuracy is achieved using only electrodes placed around the ears.
- EEG can monitor listening state and track auditory attention in two-speaker settings.

**Graphical Abstract:** EEG signals were recorded while participants listened to two concurrent speech streams, either by actively attending to one speaker or by focusing on an unrelated visual task. Spectral features of the EEG were used to classify listening state (active vs. passive) and cognitive load (low vs. high TMR). Auditory attention decoding (AAD) was performed by reconstructing the speech envelope from the EEG time signal.

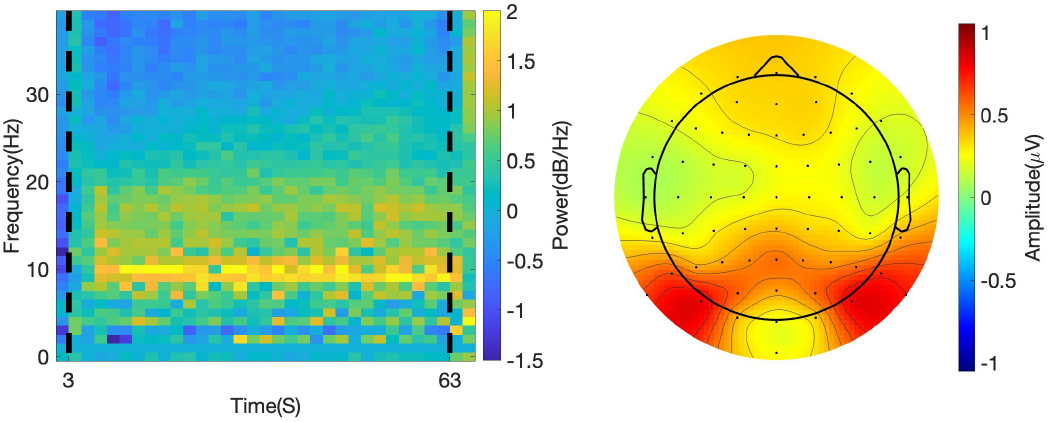

Classification of listening state (active vs. passive): 90.3% accuracy. EEG difference between active and passive listening. Left, power spectrum, right, topographic map (alpha band 8-12 Hz).

Classification of cognitive load (low vs high TMR): near chance level. EEG difference between low and high TMR. Left, power spectrum, right, topographic map (alpha band 8-12 Hz).

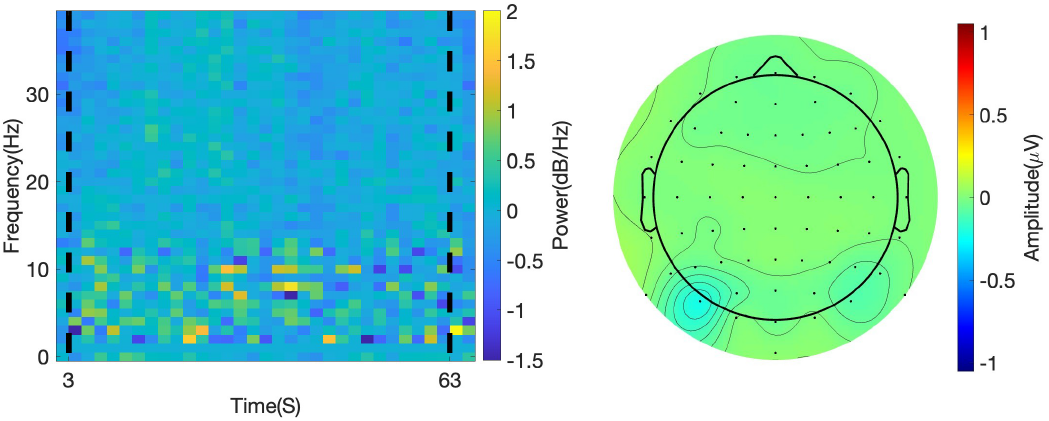

AAD achieved 84.4% accuracy, indicating robust decoding of the attended speaker during active listening, while performance dropped to near chance during passive listening.

## 1. Introduction

The cocktail-party problem—the ability to focus on a single speaker amid competing voices and background noise—is a fundamental challenge in everyday communication Cherry (1953). Successfully navigating complex acoustic scenes, like a busy restaurant or public transport, relies heavily on this skill Bizley and Cohen (2013); Shinn-Cunningham and Best (2015); Lunner et al. (2020). However, individuals with hearing impairment (HI) often find this particularly difficult, struggling to filter out irrelevant sound sources even with hearing aids Shinn-Cunningham and Best (2008); Holmes et al. (2017); Dai et al. (2018); Lunner et al. (2020); Petersen et al. (2017). These difficulties manifest as increased listening effort in noisy settings McGarrigle et al. (2014); Pichora-Fuller et al. (2016); Ohlenforst et al. (2017); Seifi Ala et al. (2020); Fiedler et al. (2021), as well as reduced cognitive aspects of listening Rönnberg et al. (2016), such as speech comprehension Peelle et al. (2011); Wendt et al. (2015) and working memory Ng et al. (2013); Arehart et al. (2013). Therefore, understanding the neural mechanisms of listening is not only relevant for auditory neuroscience, but also essential for advancing hearing technologies that aim to support users in these challenging conditions.

Electroencephalography (EEG), with its high temporal resolution Sanei and Chambers (2013); Burle et al. (2015), is widely used to monitor neural responses related to attention and auditory processing Näätänen (1990); Lalor et al. (2009); Lee et al. (2014); O’Sullivan et al. (2015). Brain-computer interface (BCI) technologies have increasingly drawn from these neural insights to build systems that can adapt to the user’s cognitive state Wolpaw et al. (2002); Curran and Stokes (2003). In the context of hearing technology, integrating principles from BCI has the potential to drive the development of hearing devices that not only amplify target sound but also decode a user’s attentional state, adapting their behavior accordingly Hill and Schölkopf (2012); O’Sullivan et al. (2015); Aličković et al. (2019); Mesgarani (2019); Ceolini et al. (2020); Han et al. (2021); Geirnaert et al. (2021); Belo et al. (2021); Hölle et al. (2021); Tanveer et al. (2024); Hjortkjær et al. (2025). Such adaptive systems could optimize sound processing strategies Lunner et al. (2020); Carlile and Keidser (2020); Slaney et al. (2020) and contribute to extended battery life by adjusting resource allocation based on detected listening states.

Distinguishing between active and passive listening states is therefore of particular interest, as detecting these states could help future hearing devices respond to the user’s attentional focus Roebben et al. (2024). Active listening involves the intentional focusing of attention on specific auditory input, while passive listening occurs without directed effort. Previous EEG studies have shown that power in the alpha (8 −12 Hz) frequency band is sensitive to attention, with lower alpha power observed during active listening compared to passive listening Dimitrijevic et al. (2017); Górska et al. (2018). However, many of these studies have used short, controlled speech stimuli, such as digit sequences Dimitrijevic et al. (2017, 2019) or texture sounds Górska et al. (2018), while everyday listening often involves continuous, complex speech in dynamic acoustic environments, where attention must be sustained over time and across competing sound sources.

At the same time, listening in noise increases cognitive load Zekveld et al. (2011), reflecting the additional cognitive resources required to process challenging auditory input, and can lead to increased listening effort Lemke and Besser (2016); Herrmann and Johnsrude (2020), which is associated with mental fatigue and reduced task performance Mattys et al. (2012); Alhanbali et al. (2019); Gagné et al. (2017); Francis and Love (2020). Listening effort is commonly described as the deliberate use of cognitive resources to overcome difficulties encountered during listening tasks Pichora-Fuller et al. (2016). Changes in alpha power have also been proposed as an objective neural marker of listening effort, with higher alpha levels observed under increased listening difficulty Seifi Ala et al. (2020); Shahsavari Baboukani et al. (2021); Fiedler et al. (2021). EEG-measured neural responses have further been identified as a physiological index capable of providing an online, continuous measure of cognitive load Antonenko et al. (2010); Schapkin et al. (2020). For hearing technology applications, monitoring these neural indicators could help identify when users experience increased effort and adjust signal processing accordingly to reduce cognitive load.

Under more realistic conditions involving continuous multi-talker speech, selective attention has been shown to modulate neural responses Mesgarani and Chang (2012); Ding and Simon (2012). Under such conditions, auditory attention decoding (AAD)— reflected by the strength of speech representation in EEG responses— has been shown to increase for target (i.e., attended) speech compared to masker (i.e., ignored) speech O’Sullivan et al. (2015); Bleichner et al. (2016); Aličković et al. (2019); Geirnaert et al. (2021); Holtze et al. (2021); Keding et al. (2024); Straetmans et al. (2024). This enhanced speech tracking for target speech has also been observed in individuals with hearing impairment Fuglsang et al. (2020); Aličković et al. (2020); Gillis et al. (2022); Keding et al. (2024); Carta et al. (2024); Wilroth et al. (2025); Tanveer et al. (2024); Sridhar et al. (2025), highlighting AAD as a robust measure of selective attention that could complement traditional spectral EEG approaches for assessing listening state and decoding attention.

Building on these insights, our study aimed to investigate EEG responses to continuous speech as indices of listening states, cognitive load, and selective attention in ecologically-valid dual-talker scenarios. Participants listened to two simultaneous talkers (male and female) presented via loudspeakers placed on the opposite side of a central screen while EEG was recorded (Figure 1). Visual tasks were presented on the central screen to divert attention from the auditory scene during passive listening. The study addressed three research questions: (1) Can listening state be decoded from neural responses, i.e., can we classify active versus passive listening? (2) How do changes in signal-to-noise ratio (SNR)—defined here as target-to-masker ratio (TMR)—affect neural responses, and can these responses be used to classify cognitive load (high vs. low TMR) during listening? (3) Can selective auditory attention be decoded from EEG signals during active listening across different TMRs?

**Figure 1:**
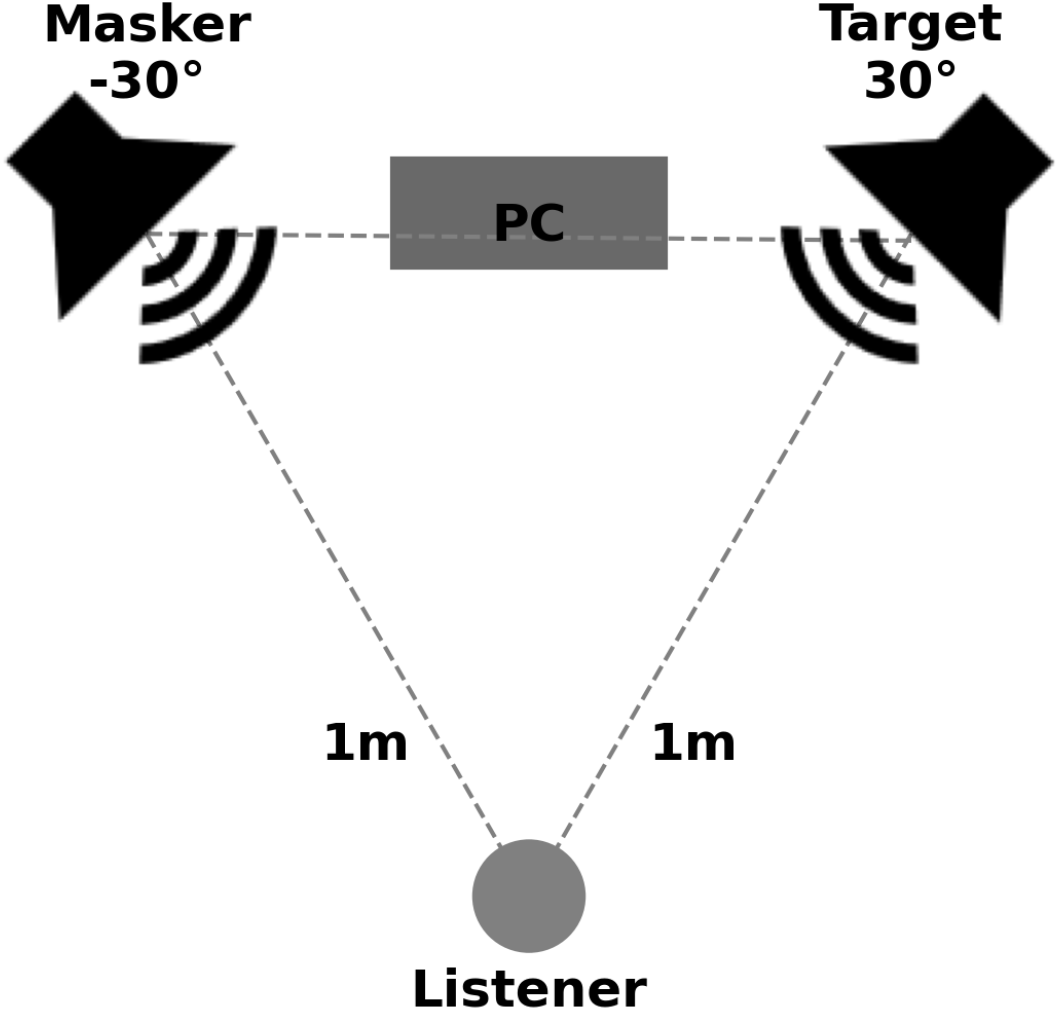
Visual and auditory stimuli delivery setup. The loudspeakers were positioned in a stereo fashion, and located 86 cm from the center of the screen.

### 2. Materials and Methods

### 2.1. Participants

The study included 15 native Danish speakers (4 females and 11 males) aged between 20 and 54, all with normal hearing. All participants had pure tone audiometric below 30 dBHL at 0.5, 1, 2 and 4 kHz.

### 2.2. Experimental Details

The study comprised 87 trials, including an initial set of 6 training trials followed by three blocks of 27 trials each. To reduce fatigue, participants were given short breaks after every eight trials and longer breaks outside the listening booth between blocks. They were seated in front of a central screen, with two loudspeakers positioned on either side (see Figure 1). Each loudspeaker played a distinct speech stream—one male and one female voice. Instructions and visual tasks were displayed on the screen. At the start of each trial, participants received on-screen instructions to either attend to one of the speech streams (from the left or right loudspeaker) or focus on the visual task.

Participants were instructed to focus on only one task at a time, either the active or passive listening task, and this instruction was repeated before the experiment to avoid multitasking. The instruction screen was displayed for 3 seconds, giving participants sufficient time to prepare for the upcoming stimuli (see Figure 2). After the instruction phase, both visual and auditory stimuli were presented for 1 minute. In the active listening tasks, two multiple-choice questions were displayed, each with three response options labeled 1 to 3. The first question addressed the general topic of the target speech, and the second referred to details from last 10 seconds of the audio. In the passive listening tasks, one multiple-choice question with 3 response options was presented. Participants indicated their response by pressing the corresponding key on the keyboard. After the response, a white screen with a central black cross was displayed for 3 seconds. This interval provided a brief pause between trials and kept the timing of the experiment consistent.

**Figure 2:**
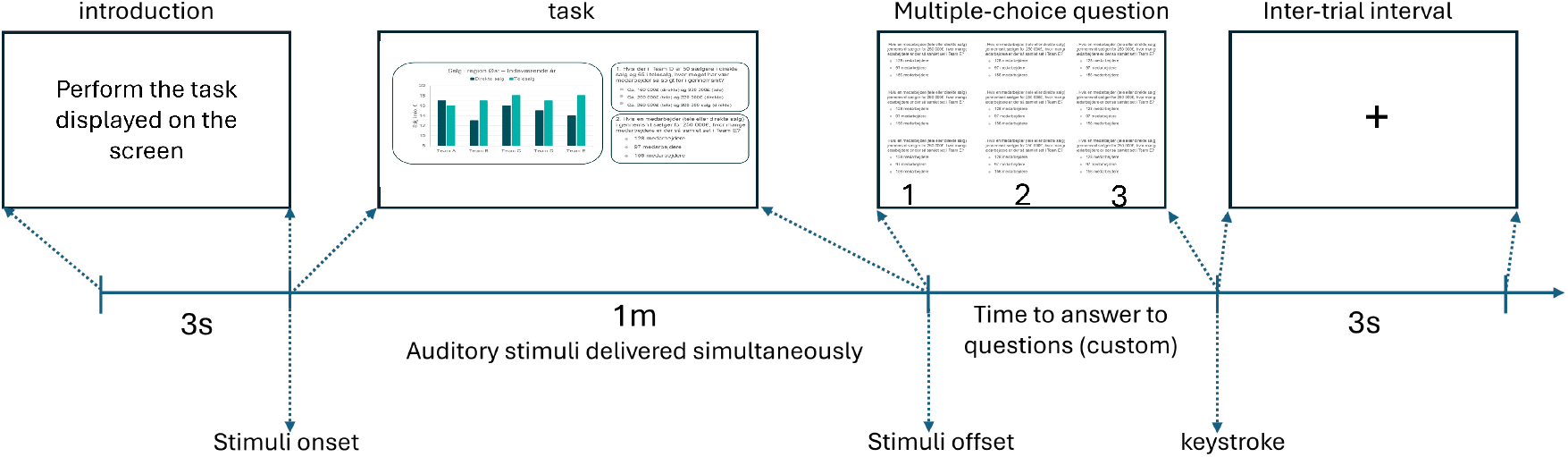
Trial Design: Participants attended to either the left or right speech stream in active listening task or the visual strimuli in passive listening task following a 3-second instruction screen. Auditory and visual stimuli were presented for 1 minute, followed by task-specific questions. A 3-second screen with a black cross separated the trials.

### 2.3. Auditory Stimuli

Auditory stimuli consisted of two simultaneous Danish news speech streams, one spoken by a male talker and the other by a female talker, each delivered from one of the loudspeaker shown in Figure 1. The target talker gender (male or female) and position (left or right) was randomized across trials, with both factors balanced throughout the experiment.

The auditory stimuli were normalized to equal root-mean-square (RMS) levels and delivered at 75 dB SPL at the listener’s ear. To assess the impact of adverse listening conditions on cognitive load, auditory tasks were presented at two distinct signal-to-noise ratio (SNR) levels, which we define here as the target-to-masker ratio (TMR), i.e., the ratio between the signal power of the target speech and the masker speech. The high SNR (TMR) condition corresponded to a +7dB difference, where the target speech was presented 7 dB above the masker (71.8 dB SPL target; 64.8 dB SPL masker). Conversely, the low SNR (TMR) condition was set at −7 dB, with the target speech 7 dB below the masker (64.8 dB SPL target; 71.8 dB SPL masker).

During trials where participants focused on the visual task, the two speech streams continued to play but were not to be actively attended. In these trials, no specific TMR was defined, as the distinction between “target” and “masker” were not applicable when the auditory task was not being performed. However, during these trials, a consistent 7 dB SPL difference between the two talkers was maintained, keeping the auditory environment similar to the listening tasks, despite participants being instructed to focus solely on the visual task.

### 2.4. Visual Stimuli

The visual stimuli included four different tasks: Spot the Difference (STD), Numerical Test (Seville), Mathematical Maze (MathMaze), and Reading Comprehension (Text). These tasks were selected to engage a range of cognitive processes, contributing to the heterogeneity of the visual condition.

#### Spot the Difference (STD)

In this task, participants were instructed to find five differences between two nearly identical images (see Figure 3), design to engage visual attention and working memory.

**Figure 3:**
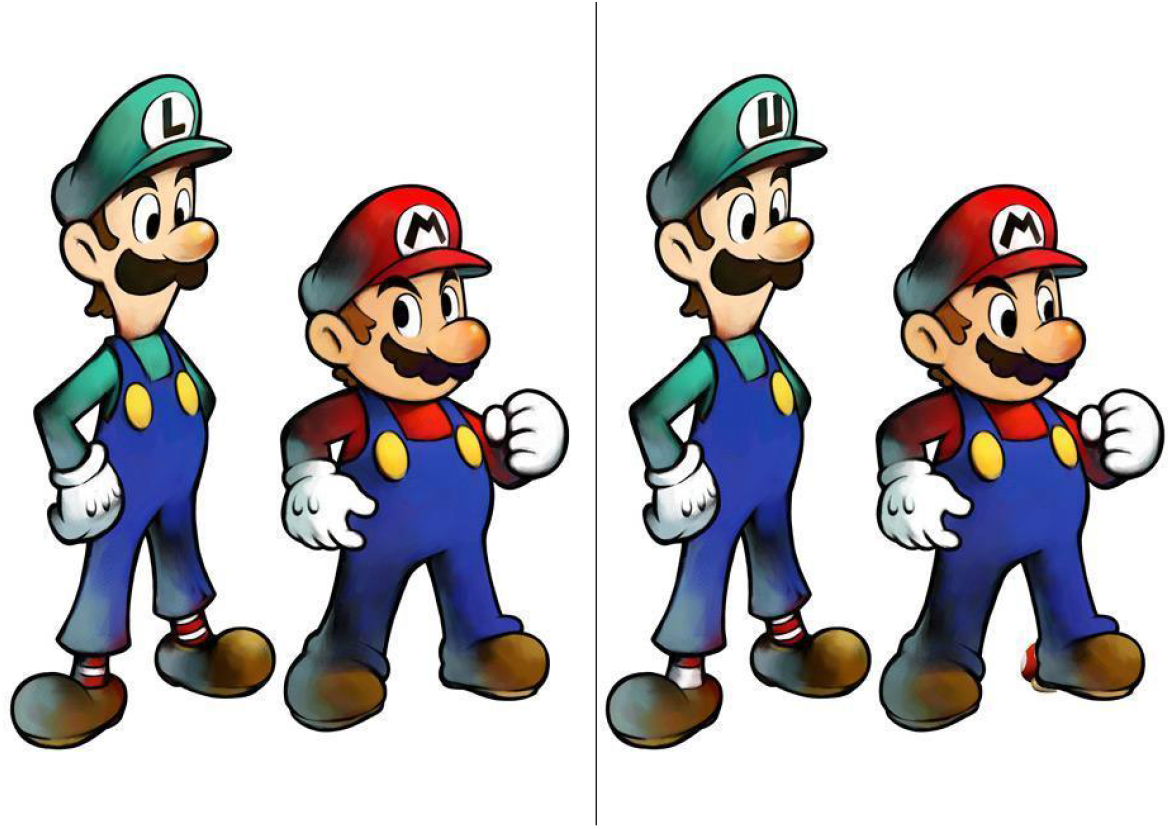
Example of the Spot the Difference (STD) task. Participants were instructed to find five differences between the two images.

#### Numerical Test (Seville)

Participants were presented with numerical information in the form of tables and graphs (see Figure 4). They answered multiple-choice questions of varying difficulty, designed to engage quantitative reasoning.

**Figure 4:**
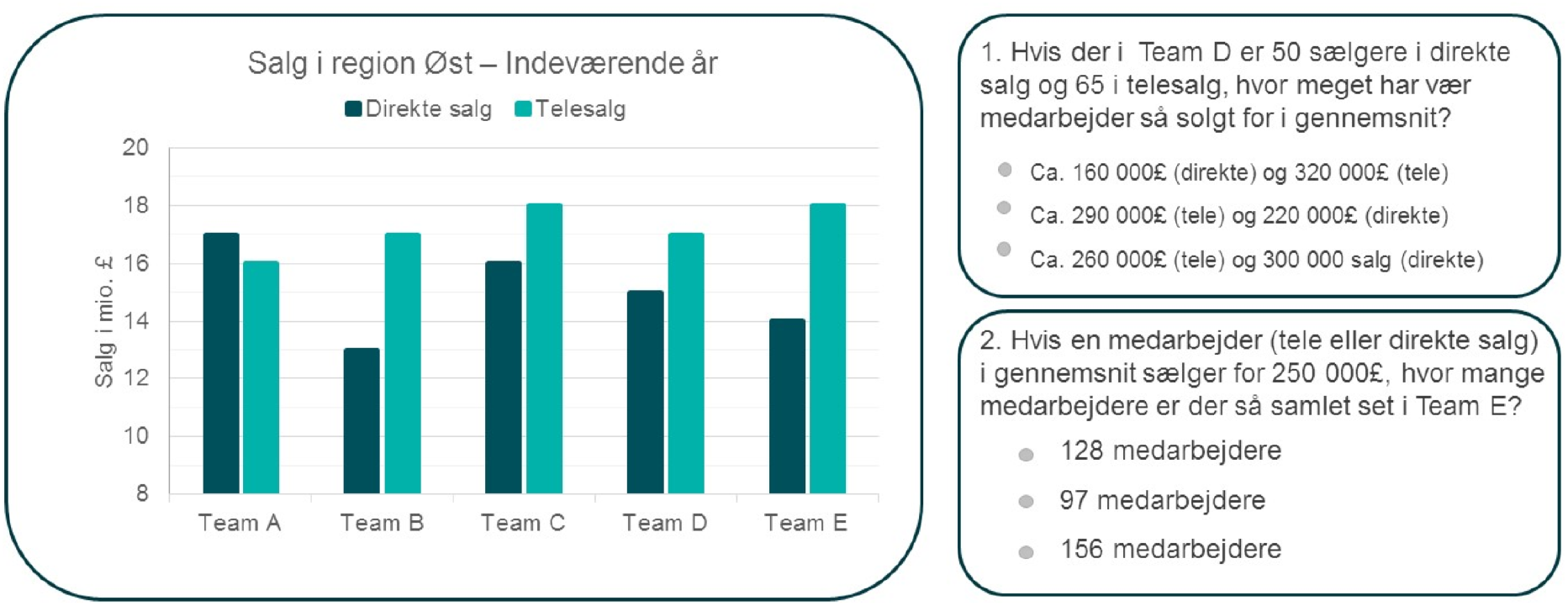
Example of the Numerical Test (Seville). Participants were instructed to answer two multiple-choice questions based on the presented diagrams.

#### Mathematical Maze (MathMaze)

Participants were instructed to find a path through a maze from the top-left to the bottom-right corner (see Figure 5). Movement was restricted to horizontal and vertical directions, and the sum of the selected numbers along the path had to equal 70. This task required logical problem-solving.

**Figure 5:**
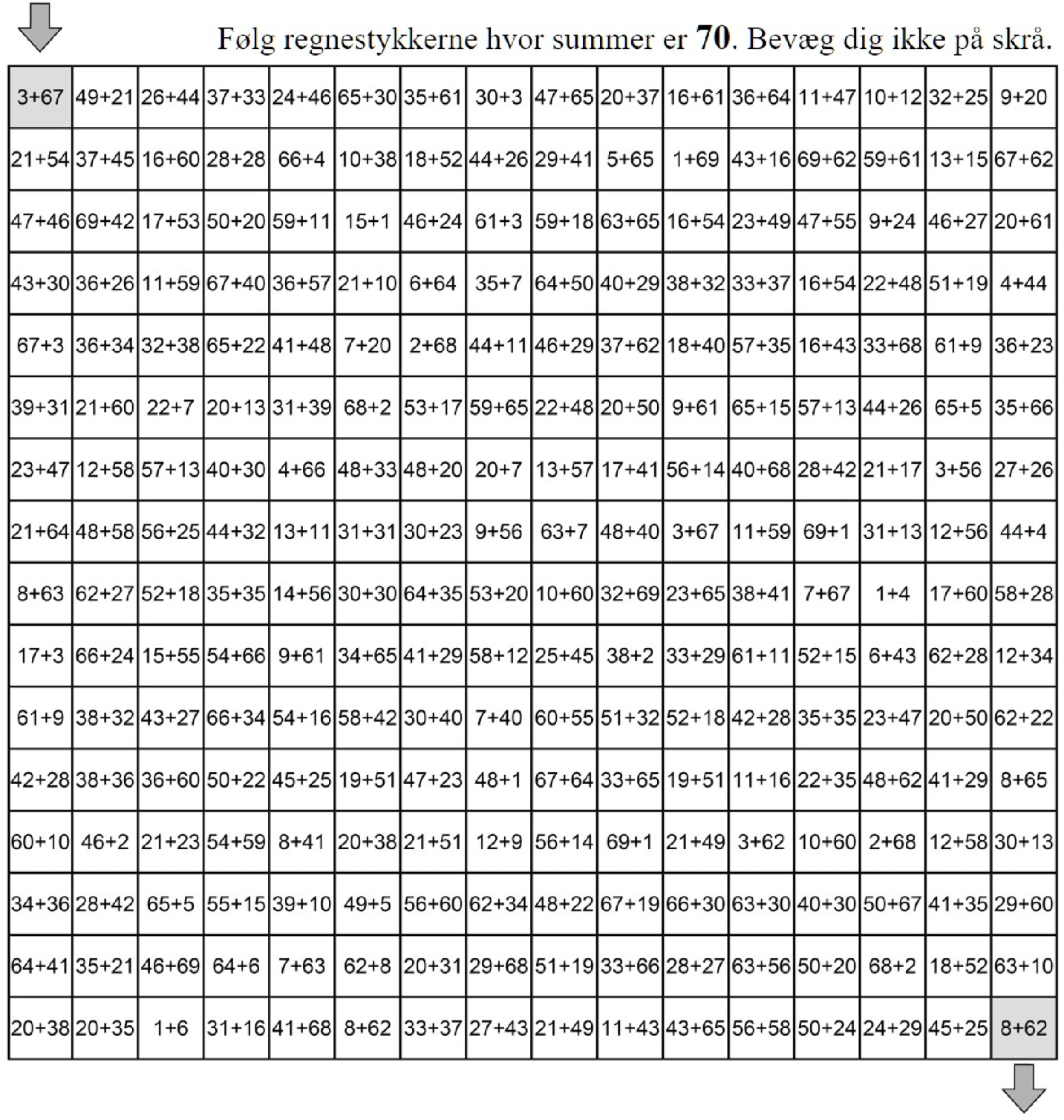
Example of the Mathematical Maze (MathMaze). Participants were instructed to navigate from the top-left corner to the bottom-right square by moving horizontally or vertically, selecting numbers that sum to 70 (instruction translated in Danish: “Find regnestykkerne hvor summen er 70”).

#### Reading Comprehension (Text)

Participants were instructed to read as much of a displayed text as possible within one minute (see Figure 6). The text covered general-interest topics drawn from sources such as Wikipedia. A comprehension question included in the text assessed understanding at a regular reading pace.

**Figure 6:**
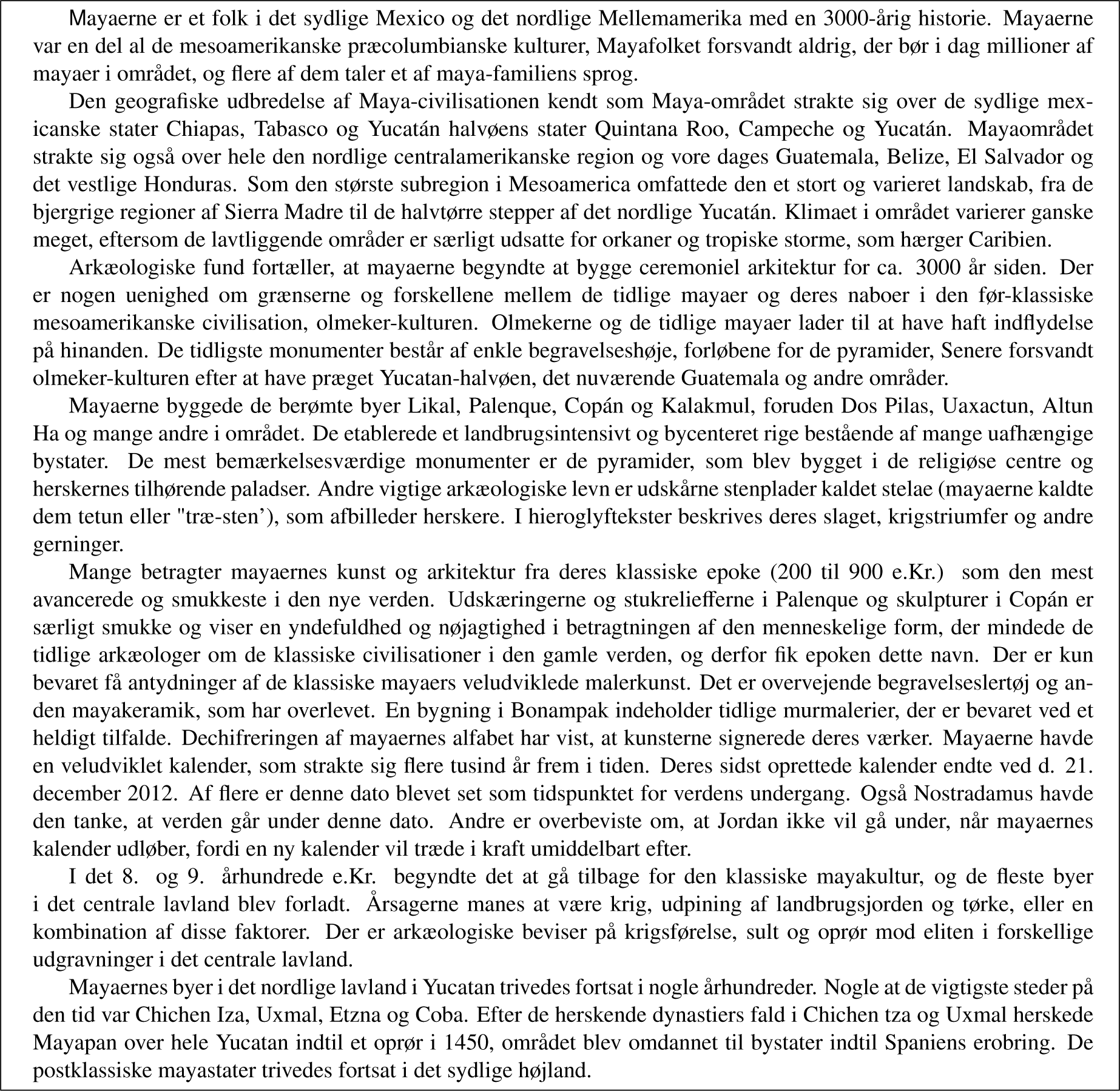
Example of the Reading Comprehension (Text) task. Participants were instructed to read as much of the text as possible within one minute.

### 2.5. Data Recordings

EEG data were recorded using 64 Ag/AgCl electrodes with a BioSemi ActiveTwo amplifier system. Two additional electrodes were placed on the mastoids to serve as offline references. The Common Mode Sense (CMS) and Driven Right Leg (DRL) electrodes provided active reference and ground, respectively. Electrode placement followed the international 10–20 system (see Figure 7). Data were acquired at a sampling rate of 1024 Hz and hardware-filtered between 0.5 and 512 Hz.

**Figure 7:**
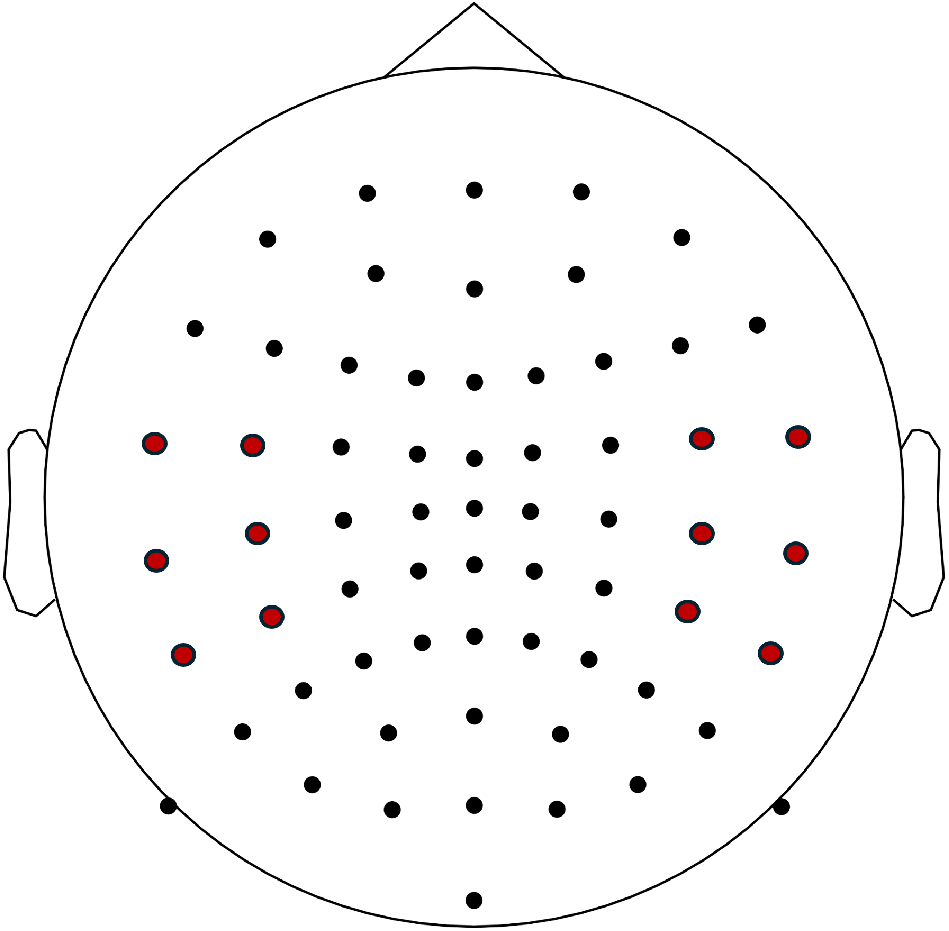
EEG electrode positions (black circles). The ear-adjacent electrodes selected for the alpha-band analysis are highlighted in red.

### 2.6. Data Pre-processing

Pre-processing steps were carried out using standard procedures to optimize signal quality and minimize artifacts:

- **Epoching:** Continuous EEG data were segmented from −3 to 63 seconds relative to stimulus onset. For analysis, only the window from 1 to 59 seconds was retained to avoid event-related potentials (ERPs) effects associated with stimulus onset and offset.
- **Event alignment:** Event co-registration was performed to synchronize EEG data with stimulus onset, allowing for precise temporal alignment of neural responses.
- **Filtering:** A 0.5 −40 Hz bandpass filter was applied to the data. A notch filter was also applied to suppress line noise.
- **Downsampling:** The signal was downsampled from 1024 Hz to 512 Hz to reduce computational demands.
- **Channel interpolation:** Channels exhibiting excessive noise or signal loss were identified by visual inspection and interpolated using neighboring channels. On average, 2.67±0.47 channels were interpolated per participant.
- **Artifact correction:** The independent component analysis (ICA) Oostenveld et al. (2011) was applied to remove components reflecting non-neural artifact (e.g., ocular or muscular activity). An average of 4.9 ± 1.2 components per participant were rejected after visual inspection.
- **Re-referencing:** Each participant’s EEG data was re-referenced to the individual common average.

## 3. Methodology

### 3.1. Spectral Power Analysis

Spectral power was quantified using the short-time Fourier transform (STFT) with a Hamming window of 1024 samples (i.e. 2 seconds) and no overlap Hayes (1996). For each time window, the power spectral density was computed via fast Fourier transform (FFT), and the energy within each frequency band -— delta (0.5 − 4 Hz), theta (4 − 8 Hz), alpha (8 − 12 Hz), and beta (12 − 25 Hz) -— was obtained by summing the squared magnitudes of the FFT bins corresponding to each band. The resulting power values were subsequently converted to dB scale 10 log10(*P*), which yielded robust estimates of power for each frequency band across all 64 EEG channels. This procedure resulted in a spectral power feature matrix of dimensions 4 × 64 features per time window.

To reduce the dimensionality of this matrix while retaining relevant neural information, univariate feature ranking was applied based on chi-square statistics Guyon and Elisseeff (2003). Each feature (i.e., band-channel combination) was evaluated for its ability to differentiate between experimental conditions (e.g., active vs. passive listening, or high vs. low TMR). From the initial set of 4 × 64 features, the 12 most discriminative features were selected for further analysis.

Complementary to this data-driven selection, a targeted feature selection was performed by manually selecting 12 electrodes located near the ears and focusing on the alpha frequency band. This choice was motivated by the potential for integrating such features into hearing aid technologies Mikkelsen et al. (2017); Lunner et al. (2020), where a limited number of strategically placed sensors may be used. The selected electrodes are indicated by red circles in Figure 7. The focus on the alpha band was guided by both prior literature and our results (see Figures 10a-10d) Seifi Ala et al. (2020); Ala et al. (2022); Shahsavari Baboukani et al. (2022); Wolf et al. (2025).

### 3.2. Classification

The spectral power feature matrix, reduced through feature selection, was used to classify listening tasks (i.e., experimental conditions). A decision tree classifier was selected for this tasks due to its interpretability and efficiency in handling non-linear feature interactions Breiman et al. (1984). The classifier was trained to distinguish between active and passive listening conditions, and within the active condition, between high and low TMR trials.

Model performance was evaluated using leave-one-out cross-validation at the trial level. For each participant, the classifier was trained on all trials except one and tested on the held-out trial. This procedure was repeated for all trials, ensuring that classification performance was assessed on data not used during training. This within-subject validation approach provides a reliable estimate of the model’s generalization performance across conditions.

### 3.3. Auditory attention decoding (AAD)

AAD was performed using a Stimuli Reconstruction approach, in which the goal is to reconstruct the speech envelope *S* from EEG recodings *R* using a linear decoder *g* O’Sullivan et al. (2015); Aličković et al. (2019); Geirnaert et al. (2021); Crosse et al. (2021). This backward model allows us to assess how well the neural responses track the target speech stream.

The auditory stimuli, including target and masker signals, were first resampled to 44, 096 Hz, a multiple of 64 Hz. Speech envelopes were extracted using the Hilbert transform, followed by downsampling to 64 Hz and bandpass filtering between 1− 8 Hz using a sixth-order Butterworth filter. EEG signals were preprocessed in the same way to ensure temporal alignment with the speech envelopes.

In the test phase, the reconstructed speech envelope *Ŝ*_*test*_ test was computed as:

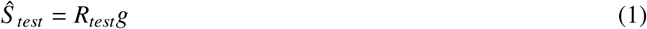

The decoder *g* was estimated during training by minimizing the mean-squared error between the original and reconstructed speech envelopes. The solution to this least-squares optimization problem, with ridge regularization, is given by Aličković et al. (2019); Geirnaert et al. (2021):

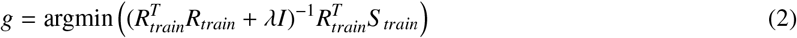

Here, λ is the regularization parameter Wong et al. (2018) that was tuned from the range 10−6 to 106 to improve reconstruction performance.

The model was estimated using target speech by leave-one-out cross-validation approach, where all trials except one were used for estimating each participant’s decoder *g*. Since participants were presented with two simultaneous speech streams, two stimuli can be reconstructed per trials. The reconstruction accuracy was assessed as the Pearson’s *r* correlation coefficient between the reconstructed *Ŝ*_*test*_ and the original *S*_*test*_ speech envelopes. Decoding accuracy was calculated as the proportion of trials in which the correlation was higher for the target speech compared to the masker speech.

Consistent with the power analysis, a subset of 12 electrodes near the ears was manually selected for stimulus reconstruction.

### 4. Statistical Analysis

To investigate differences in performance and brain activity related to listening task (active vs. passive listening) and listening condition (low vs. high TMR), the following statistical methods were used:

- **Wilcoxon Rank-Sum Test**: Used to compare differences between the active and passive listening tasks in terms of response accuracy. This test was also applied to assess statistical differences between active and passive listening tasks in the time-frequency domain of EEG signal. Results are visualized as figures showing p-values across time and frequency bins.
- **One-Way Analysis of Variance (ANOVA)**: Applied to evaluate variations in response accuracy across the different visual task types (text, spot-the-difference, math maze, and Seville tasks), as well as to explore differences in spectral power in passive listening task. Post-hoc analyses were conducted to identify pairwise differences between task types where significant effects were observed.

## Results

### 5.1. Decoding listening state (active vs. passive listening)

Neural data revealed clear differences between active and passive listening. Spectrograms averaged over EEG channels, trials, and participants illustrate differences between active and passive listening conditions (Figures 8a, 8b, and 8c) . These figures reveal increased power during speech presentation (3 to 63 seconds) in the active listening condition relative to the passive listening, particularly within the alpha frequency bands. These findings are supported by statistical analyses (Figure 8d), where the Wilcoxon Rank-Sum test revealed significant differences (low p-values) in the alpha band during speech presentation.

**Figure 8:**
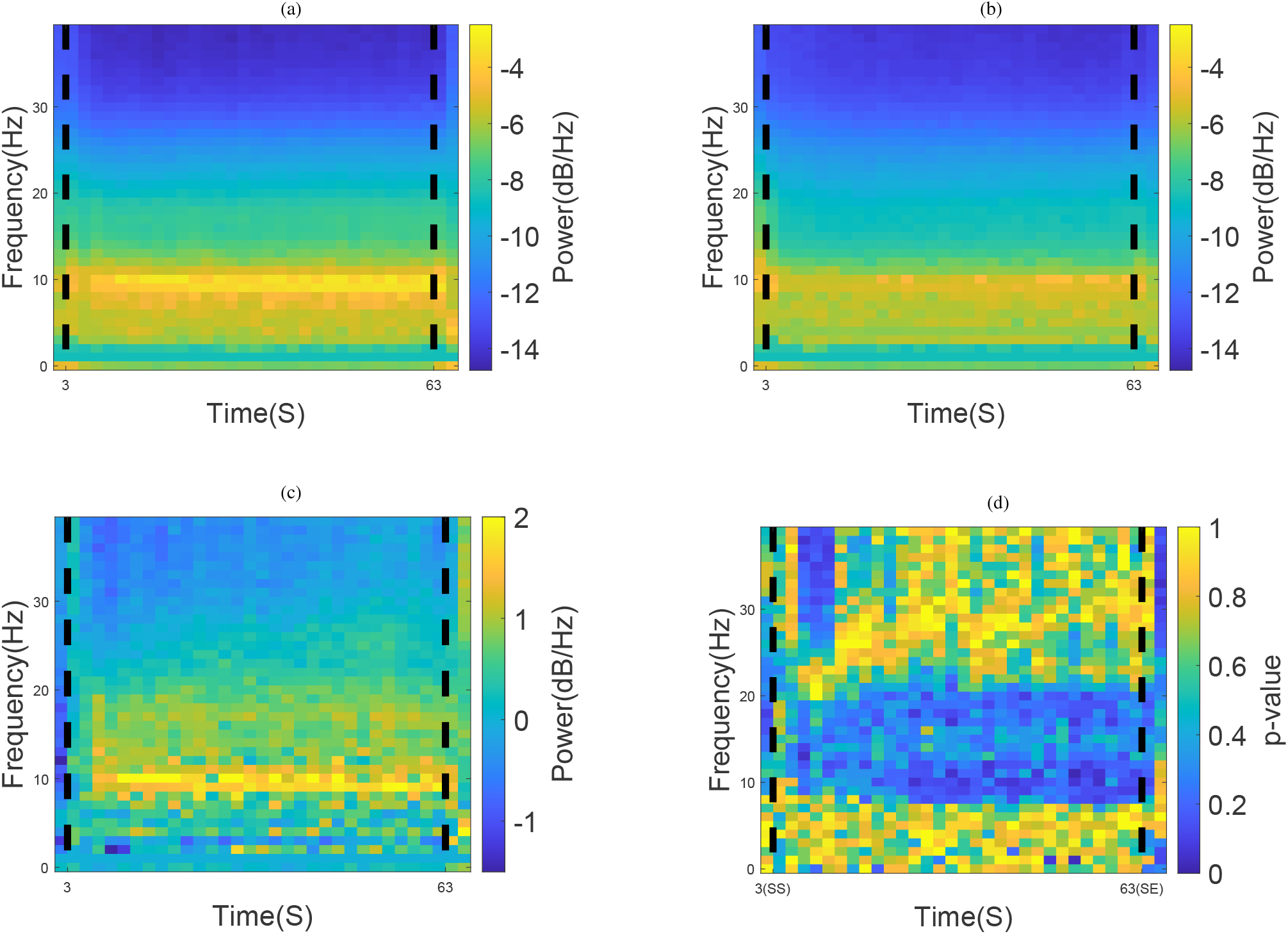
Spectrogram overviews averaged over channels, trials, and participants, for (a) active listening, (b) passive listening and (c) active - passive difference. Statistical test results displaying p-values for (d) ranksum test of active vs. passive listening.

EEG topographic maps averaged across participants, trials, and the stimulus window (3 − 63 seconds) further highlighted these differences. Active listening elicited stronger alpha activity, particularly in the parietal regions, compared to passive listening (Figures 9a, 9b, and 9c).

**Figure 9:**
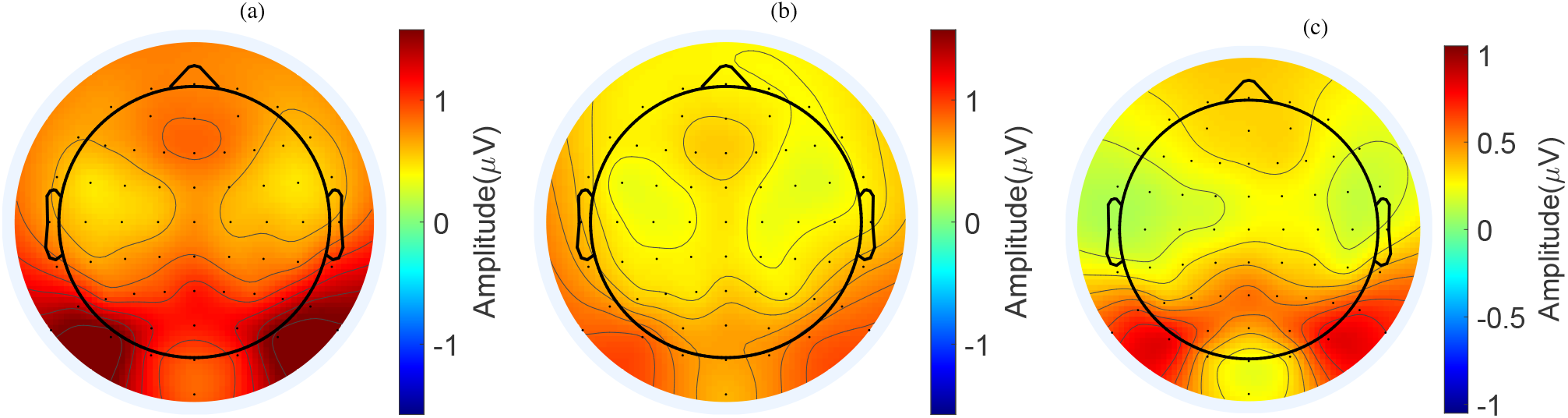
EEG topographic maps averaged across participants, trials, stimuli window (3 −63s), and the alpha band (8 − 12 Hz). Panels show (a) active listening, (b) passive listening, (c) active - passive difference.

Figure 10 shows the absolute log-transformed p-values from the univariate feature ranking analysis across the four frequency bands, which were used as the basis for the classification of active versus passive listening. Averaged across all participants, the results indicate that features in the alpha band yielded the highest absolute log-p values, reflecting lower p-values and greater informativeness.

**Figure 10:**
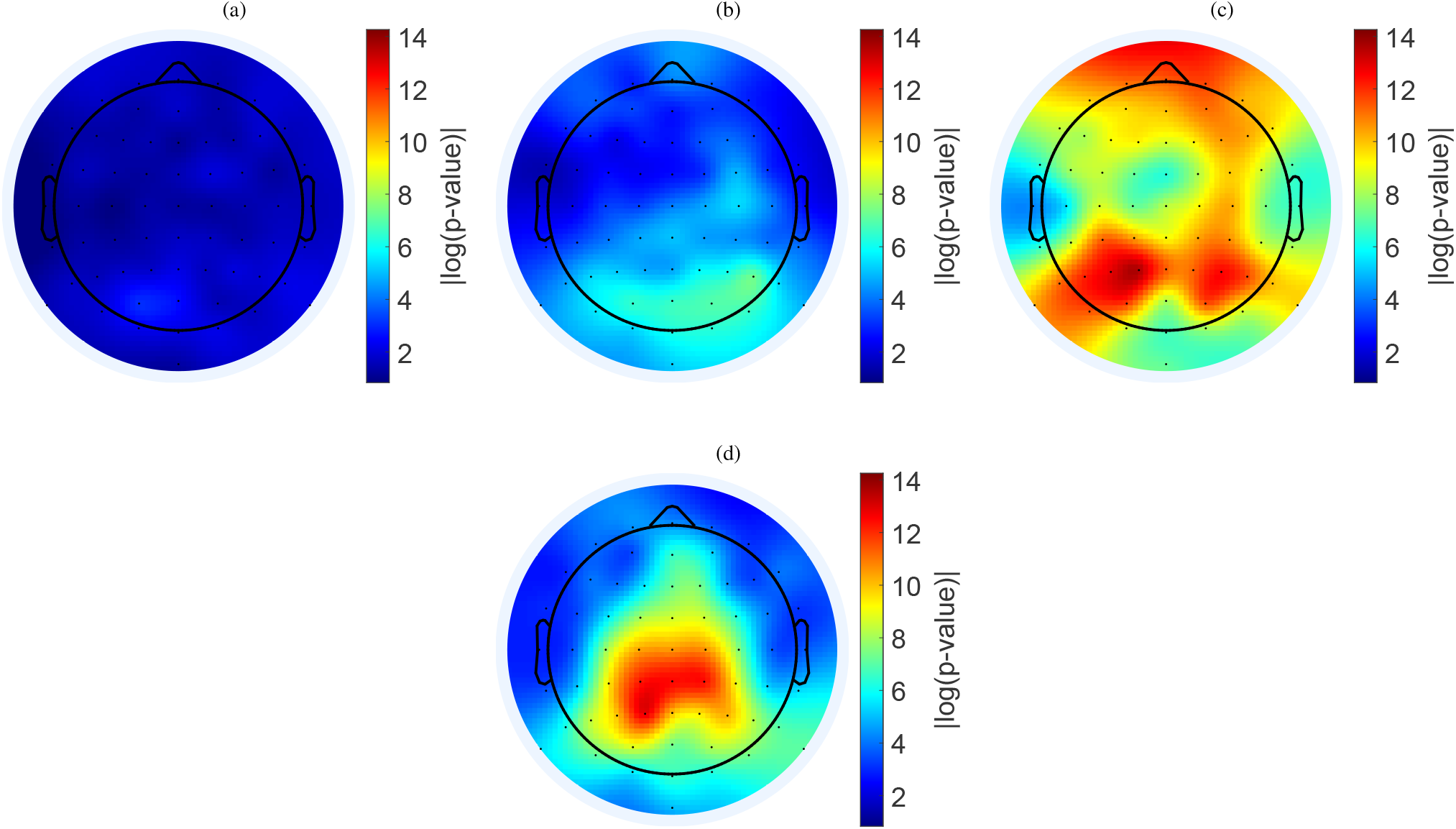
Feature selection results showing log-transformed p-values from univariate feature ranking analysis across four frequency bands: (a) delta (0.5-4 Hz), (b) theta (4-8 Hz), (c) alpha (8-12 Hz), and (d) beta (12-25 Hz).

Using a decision tree classifier with the best 12 features selected from all EEG channels and frequency bands, classification between active and passive listening achieved high accuracy across participants, ranging from 77.8% to 98.8%, with an average accuracy of 90.3% (Table 1, first row). Limiting the analysis to a subset of electrodes around the ears resulted in classification accuracy between, 64.2 and 97.5, with an average accuracy of 81.9% (Table 1, second row). This suggests that features derived from these specific electrodes are still sufficiently informative for distinguishing active from passive listening.

**Table 1.**
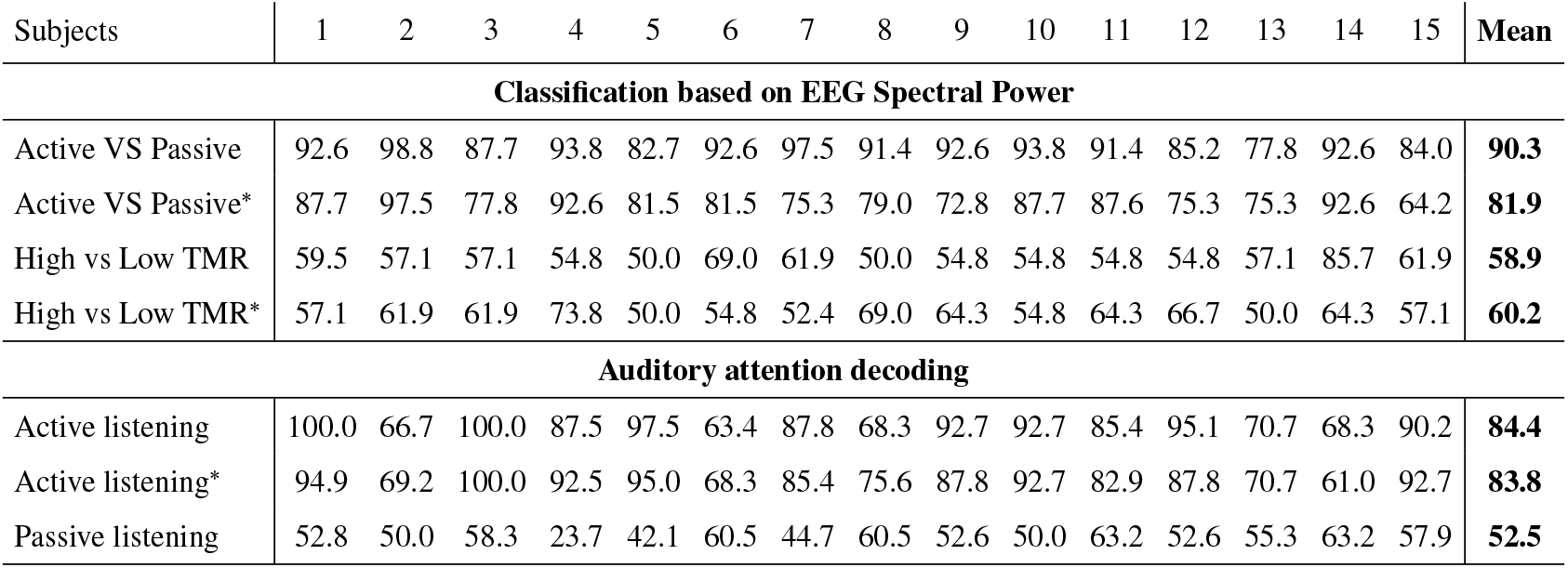
Classification results based on EEG spectral power (top) and auditory attention decoding (AAD) (bottom). The * denotes the analysis done using only around the ear electrodes, shown in red coulour in Figure 7.

| Subjects | 1 | 2 | 3 | 4 | 5 | 6 | 7 | 8 | 9 | 10 | 11 | 12 | 13 | 14 | 15 | Mean |
| --- | --- | --- | --- | --- | --- | --- | --- | --- | --- | --- | --- | --- | --- | --- | --- | --- |
| **Classification based on EEG Spectral Power** |  |  |  |  |  |  |  |  |  |  |  |  |  |  |  |  |
| Active VS Passive | 92.6 | 98.8 | 87.7 | 93.8 | 82.7 | 92.6 | 97.5 | 91.4 | 92.6 | 93.8 | 91.4 | 85.2 | 77.8 | 92.6 | 84.0 | **90.3** |
| Active VS Passive* | 87.7 | 97.5 | 77.8 | 92.6 | 81.5 | 81.5 | 75.3 | 79.0 | 72.8 | 87.7 | 87.6 | 75.3 | 75.3 | 92.6 | 64.2 | **81.9** |
| High vs Low TMR | 59.5 | 57.1 | 57.1 | 54.8 | 50.0 | 69.0 | 61.9 | 50.0 | 54.8 | 54.8 | 54.8 | 54.8 | 57.1 | 85.7 | 61.9 | **58.9** |
| High vs Low TMR* | 57.1 | 61.9 | 61.9 | 73.8 | 50.0 | 54.8 | 52.4 | 69.0 | 64.3 | 54.8 | 64.3 | 66.7 | 50.0 | 64.3 | 57.1 | **60.2** |
| **Auditory attention decoding** |  |  |  |  |  |  |  |  |  |  |  |  |  |  |  |  |
| Active listening | 100.0 | 66.7 | 100.0 | 87.5 | 97.5 | 63.4 | 87.8 | 68.3 | 92.7 | 92.7 | 85.4 | 95.1 | 70.7 | 68.3 | 90.2 | **84.4** |
| Active listening* | 94.9 | 69.2 | 100.0 | 92.5 | 95.0 | 68.3 | 85.4 | 75.6 | 87.8 | 92.7 | 82.9 | 87.8 | 70.7 | 61.0 | 92.7 | **83.8** |
| Passive listening | 52.8 | 50.0 | 58.3 | 23.7 | 42.1 | 60.5 | 44.7 | 60.5 | 52.6 | 50.0 | 63.2 | 52.6 | 55.3 | 63.2 | 57.9 | **52.5** |

### 5.2. Classification of cognitive load (low vs. high TMR)

Task performance was assessed using responses to questions at the end of each trial (Figures 11 and 12). Average accuracy across participants consistently exceeded 80% across all conditions, including high and low TMRs and the different visual tasks. Statistical comparison revealed no significant difference in accuracy between high and low TMR conditions (p = 0.38), indicating that variations in acoustic listening difficulty did not substantially affect participants’ ability to answer questions correctly.

**Figure 11:**
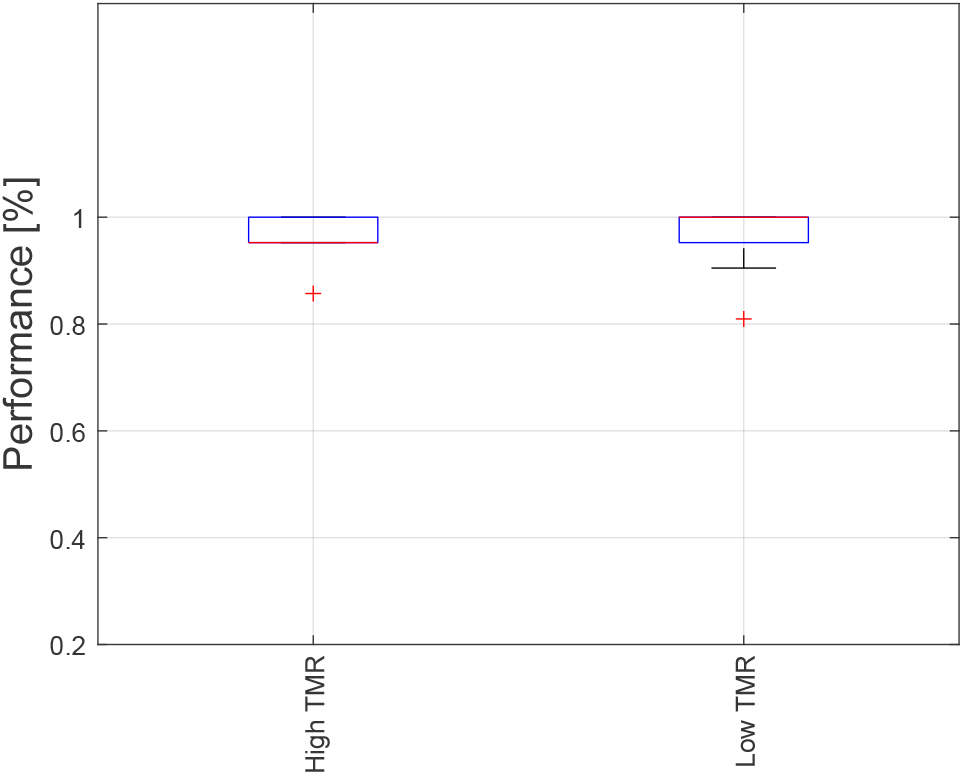
Performance analysis results, measured as the ratio of correctly answered multiple-choice questions, illustrating the average performance across trials and participants for the Low TMR vs High TMR conditions.

**Figure 12:**
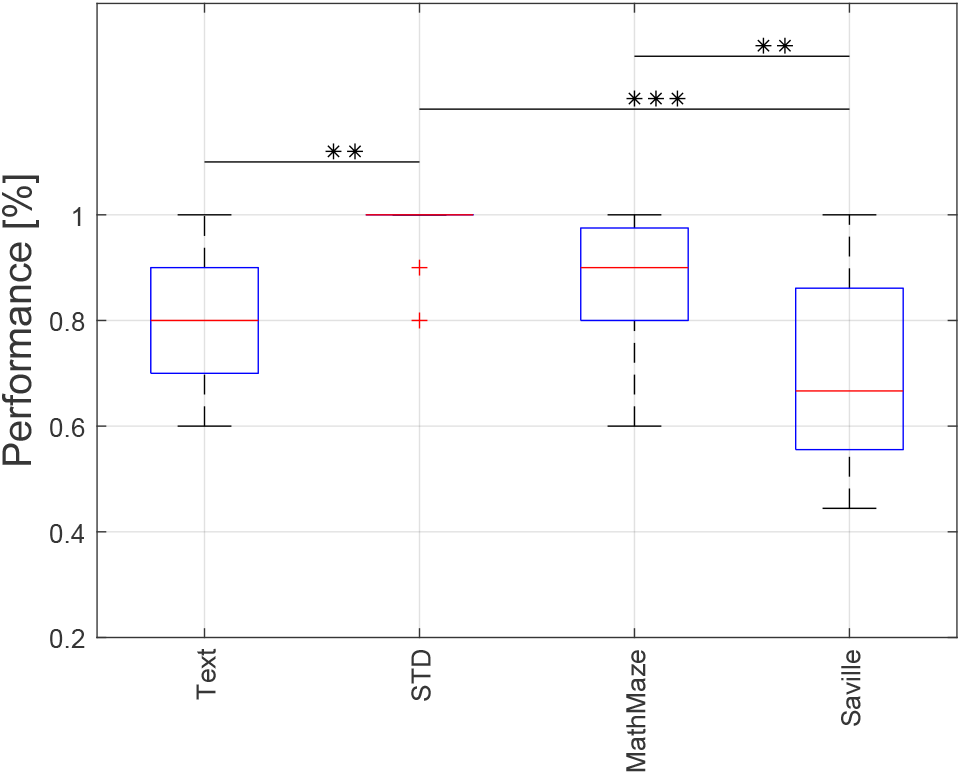
Performance analysis results, measured as the ratio of correctly answered multiple-choice questions, illustrating the average performance across trials and participants for the different visual tasks. Posthoc analysis of ANOVA test between four visuals stimuli are also shown using strikes.* indicates p-values less than 0.05, ** indicates p-values less than 0.001, and *** indicates p-values less than 0.0001

In contrast, the type of concurrent visual task significantly influenced performance. A one-way ANOVA (*p* ≤0.0001) revealed differences in accuracy across the Text, Spot-the-Difference (STD), MathMaze, and Seville tasks. Post-hoc analysis (Figure 12) identified significant differences between the Text and STD tasks, as well as between STD and both MathMaze and Seville tasks. These results suggest that cognitive load or engagement associated with the visual tasks had a measurable effect on task performance, independent of TMR.

Neural measures were consistent with the behavioral findings. Spectrograms averaged across EEG channels, trials, and participants showed minimal differences between high and low TMR conditions (Figure 13a, 13b, and 13c). Statistical analyses confirmed this, with high p-values across nearly all time-frequency bins 13d, indicating no significant difference in spectral power. Similarly, EEG topographic maps in the alpha band (Figures 14a, 14b, and 14c) revealed highly similar spatial distributions for high and low TMR conditions, further supporting the absence of detectable neural effects.

**Figure 13:**
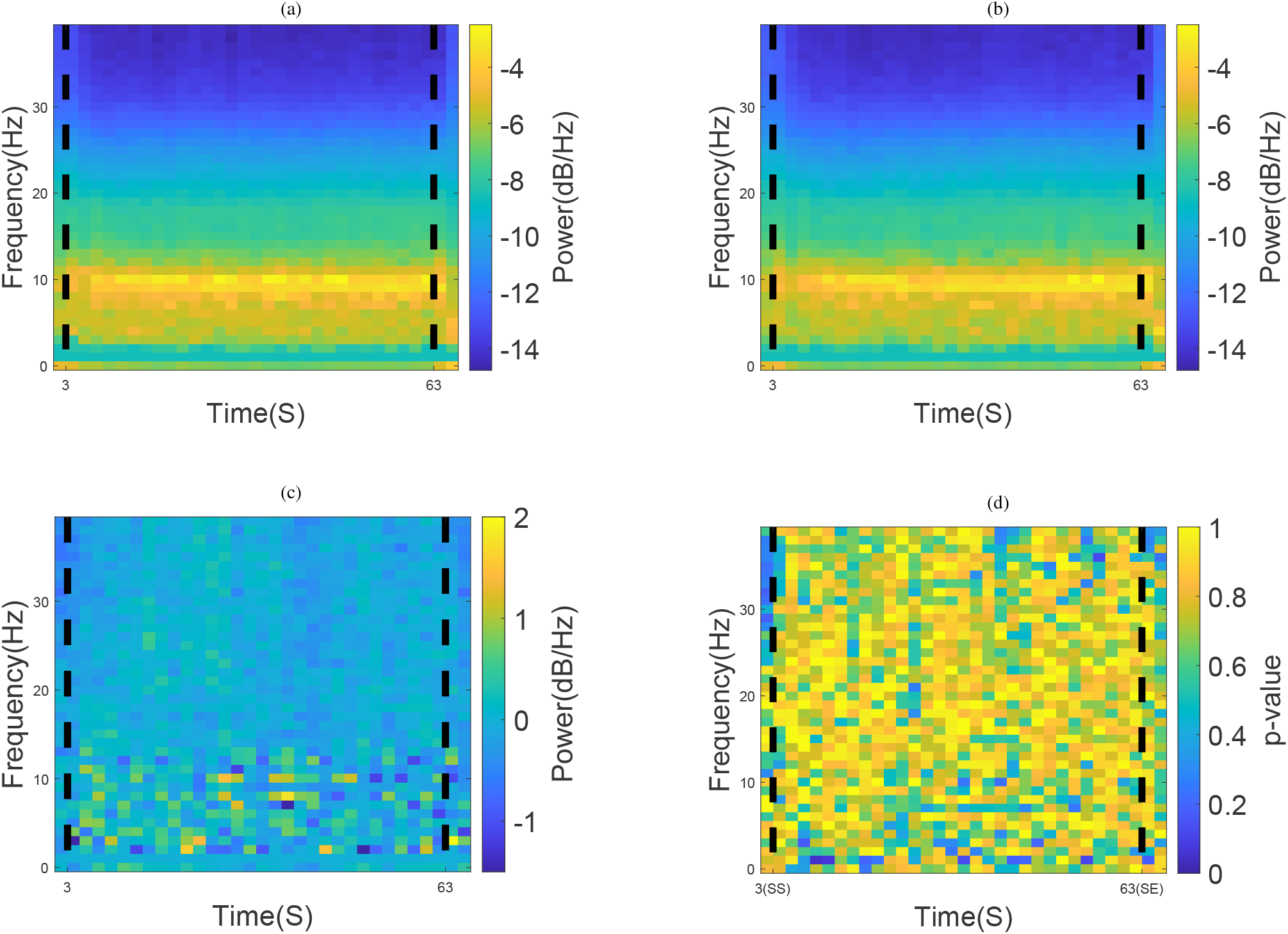
Spectrogram overviews averaged over channels, trials, and participants, for High TMR (a) and Low TMR (b), High - Low TMR difference (c). Statistical test results displaying p-values for ranksum test of High vs. Low TMR (d).

**Figure 14:**
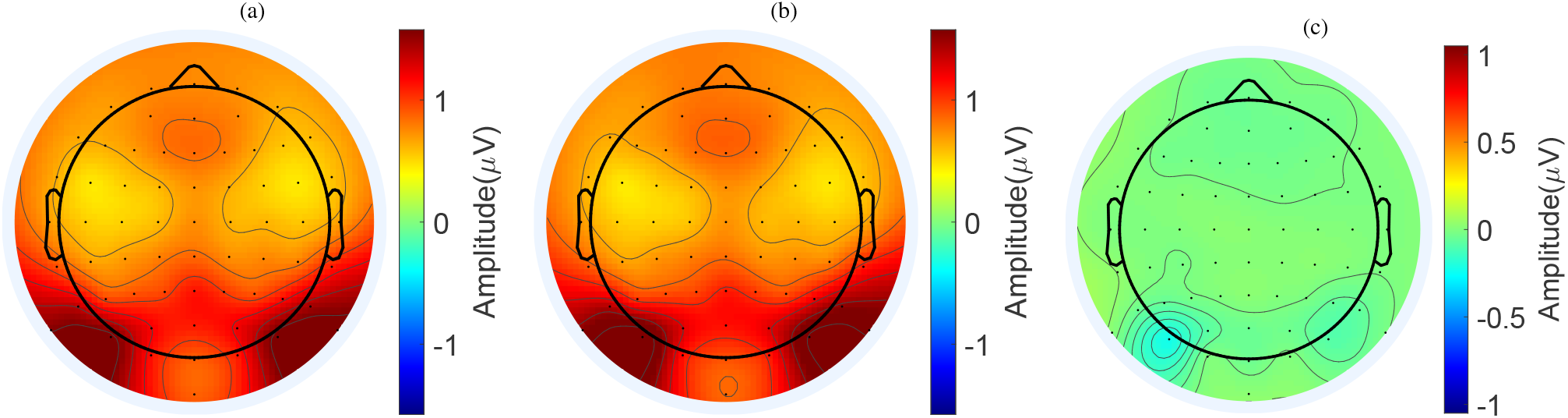
EEG topographic maps averaged across participants, trials, stimulus window (3 −63s), and alpha band (8 −12 Hz). Panels show (a) high TMR, (b) low TMR and (c) High - Low TMR difference.

Classification analyses reinforced these observations. Attempts to distinguish high versus low TMR conditions based on EEG spectral features resulted in near-chance performance, with average accuracies of 58.9% across all electrodes and 60.16% using only electrodes around the ears (Table 1, third and forth rows). These results suggest that EEG spectral power alone was insufficient to differentiate TMR conditions. This lack of a neural effect, combined with the behavioral data, suggests that the acoustic manipulations were not challenging enough to induce a measurable difference in cognitive load within the tested range.

Overall, both behavioral and neural data indicate that, within the tested TMR range, acoustic listening difficulty did not significantly modulate cognitive load. By contrast, differences in visual task demands had a clear impact on performance.

### 5.3. Auditory attention decoding (target vs. masker)

To evaluate whether neural data could distinguish attended (target) versus unattended (masker) speech, stimulus reconstruction models were trained using EEG data as described in section 3.3. During active listening, models trained on the target speech achieved an average classification accuracy of 84.42% across participants (Table 1, fifth row). Using only electrodes around the ears maintained high accuracy (83.8%, Table 1, sixth row).

In contrast, during passive listening—when participants attended to visual stimuli rather than speech—classification accuracy using all electrodes dropped to 52.5% (Table 1, seventh row). This near-chance performance confirms that AAD relies on active attention, demonstrating that the method can reliably distinguish target from masker speech during active listening but not during passive listening when attention is directed elsewhere.

### 6. Discussion

This study investigated the neural dynamics associated with active and passive listening using long continuous speech. It also examined the feasibility of EEG-based features for assessing cognitive load and decoding auditory attention during active listening. By using ecologically valid stimuli, the study provides insights into how neural responses reflect real-world listening demands, which could inform future applications in hearing aid technologies.

Our results demonstrate high classification accuracy for distinguishing active versus passive listening, with an average accuracy of 90.3% using all EEG channels. The finding that a similarly high accuracy of 81.9% was achieved with only electrodes around the ears is particularly significant. This highlights the potential of using more practical and unobtrusive electrode configurations for future applications, such as hearing aids Ala et al. (2022); Roebben et al. (2024).

In contrast, classification of cognitive load based on high versus low TMR conditions showed near-chance performance (58.9%), even when considering only electrodes around the ears (60.16%). One likely explanation is that the TMR difference (7 dB) was not large enough to elicit detectable changes in cognitive demands, resulting in largely similar EEG signals. This is consistent with the role of binaural masking release Sutojo et al. (2020), where spatially separated sound sources reduce energetic masking compared to co-located sources. Future studies could increase listening difficulty by introducing more competing voices, reducing spatial separation between sources, or by employing an adaptive speech reception threshold (SRT) procedure to more precisely control individual listening difficulty levels.

Beyond TMR, other factors could also have influenced our findings. Voice similarity, for instance, has been shown to be major factor affecting cognitive processing load during listening, often outweighing the effects of spatial separation between target and masker speech Zekveld et al. (2014). In the present study, voices had similar characteristic tempo and pronunciation, as they were both from Danish news broadcasts. However, the male and female voices had a clear frequency difference, which may have contributed to the lack of neural differences observed between high and low TMR condition. Additionally, it would also be valuable to study populations with different hearing abilities, as cognitive load is known to affect speech motor performance differently in older versus younger adults MacPherson (2019), and that even low background noise increases cognitive load in older adults listening to competing speech Meister et al. (2018). Future work should also consider explicitly incorporating individual cognitive factors, which are known to modulate how listening difficulty is reflected in neural activity.

Finally, the analysis of target versus masker speech highlights the potential of EEG-based BCIs for decoding auditory attention. During active listening, AAD accuracy was relatively high (84.42%), indicating that EEG responses can effectively track a listener’s attention. This finding is in line with previous research O’Sullivan et al. (2015); Wong et al. (2018); Aličković et al. (2019); Mesgarani (2019); Aroudi and Doclo (2020); Geirnaert et al. (2021); Straetmans et al. (2022, 2024); Hjortkjær et al. (2025), which has shown significant decoding accuracy when identifying target speech, even under challenging SNR conditions Schäfer et al. (2018); Das et al. (2018) and across different populations, including listeners with hearing impairment Fuglsang et al. (2020); Aličković et al. (2020, 2021); Sridhar et al. (2025); Wilroth et al. (2025). Comparable accuracy (83.8%) was also obtained using only electrodes positioned around the ears, in line with recent work on ear-centered EEG Bleichner et al. (2016); Mirkovic et al. (2016); Holtze et al. (2022); Borges et al. (2024); Thornton et al. (2025); Geirnaert et al. (2025). This suggests that practical and unobtrusive electrode configurations may be sufficient for future real-world applications.

The observed low decoding accuracy (52.5%) in decoding the target talker in the passive listening condition aligns with the experimental design, where participants were explicitly instructed to focus on the visual task and neglect the auditory stimuli. This results in a lack of neural correlates associated with selective attention. Therefore, the decoder faces challenges due to the lack of attention to the corresponding auditory information, leading to performance around chance level.

## 7. Conclusion

This study demonstrates that EEG signals can reliably distinguish between active and passive listening and decode a listener’s selective auditory attention during continuous speech. The successful classification using a limited set of ear-centered electrodes underscores the significant potential for practical, real-world applications in hearing aids and auditory brain-computer interfaces (BCIs). In contrast, our attempt to classify cognitive load based on a 7 dB TMR difference yielded near-chance performance, which highlights a key challenge: the acoustic manipulation was insufficient to elicit a robust neural response reflecting a change in listening effort. Future work should investigate stronger variations in listening difficulty, include populations with diverse hearing abilities, and consider factors such as voice similarity. In summary, this study confirms the potential of EEG-based BCIs to monitor a user’s attentional state and offers a foundation for developing objective measures of listening and next-generation hearing devices that dynamically adapt to a listener’s cognitive state.

## Acknowledgments

The authors acknowledge Thomas Lunner for his contributions to the design of the experiment, and Eline Borch Petersen for her contributions to the experimental design, setup, and data acquisition.

